# Tolerance of Wildlife in Protected Area Borderlands

**DOI:** 10.1101/2021.03.27.436188

**Authors:** Leandra Merz, Elizabeth F. Pienaar, Timothy Fik, Shylock Muyengwa

## Abstract

Increases in human-wildlife conflict globally threaten human wellbeing and biodiversity conservation. Sustainable solutions that promote coexistence of people and wildlife are needed, especially in human-dominated landscapes surrounding protected areas. People’s attitudes toward wildlife influence their behaviours, including tolerance for human-wildlife interactions, poaching, and habitat degradation. Better understanding of how to improve people’s attitudes toward wildlife is instrumental to promoting coexistence between people and wildlife in shared spaces. Efforts to promote coexistence often fail because they are based on the inaccurate assumption that people’s attitudes towards wildlife are directly and proportionally related to wildlife-based financial incentives and costs. In reality, people’s attitudes towards wildlife are far more complex. We analysed surveys (n=237) from Mozambique to examine people’s attitudes toward wildlife in the buffer zones surrounding protected areas using logistic regression and Getis-Ord hot-spot analysis (GI*). Mozambique, which is under-represented in the wildlife-based research literature, is characterized by extreme poverty and rewilding efforts. We found that most respondents were tolerant of wildlife and tolerance was positively correlated with people’s age, gender, and agreement with rules governing wildlife conservation. People’s tolerance for wildlife was also reinforced if they receive benefits from wildlife and are situated further from the park fence. Predation, human harm, and crop loss were not significant predictors of tolerance. We found no evidence of spatial patterns in tolerance for wildlife. Our results suggest that wildlife conservation programs are more likely to be successful if benefits are distributed equitably and community members are actively involved in decision making.

## Introduction

With the human population expanding globally, there is increased encroachment of humans into landscapes that were previously dominated by wildlife (Fischer et al., 2011). This encroachment leads to more encounters between humans and wildlife and consequently increased frequency and intensity of conflict (Newmark et al., 1993; Barua et al., 2013). Human-wildlife conflicts encompass interactions between humans and wildlife that negatively affect one or both groups (Thorn et al., 2012) and conflicts between people over how to manage wildlife (Kansky et al., 2016). Conflicts are predicted to increase as the human population grows and demands more resources (Kansky & Knight, 2014), including access to land. This is problematic because protected areas cover less than 13% of the globe and most of the world’s protected areas are too small to support sustainable populations of large mammals that have extensive home range requirements (Newmark et al., 1993; Barua et al., 2013). Many species rely on privately and communally owned land outside protected areas as part of their home ranges or as corridors that provide connectivity to other protected areas. Reliance on land outside of protected areas brings wildlife into direct contact with humans who utilize this land for cattle grazing, farming, and other activities. Given the limited availability of land and money for conservation, it is vital to develop strategies that will allow humans and wildlife to coexist spatially and temporally, especially on land bordering protected areas (Thorn et al., 2012).

Efforts to promote coexistence between people and wildlife are challenging because of the multi-faceted and interconnected drivers and impacts of conflict. High population growth rates and resulting increases in human-wildlife conflicts are a major threat to wildlife conservation and human lives and livelihoods in sub-Saharan Africa (Nicole, 2019). Human-wildlife conflict in sub-Saharan Africa frequently manifests as wildlife destroying crops and property, wildlife preying on livestock, wildlife injuring or killing humans, the transmission of pathogens between wildlife and livestock, people killing wildlife, and people destroying or degrading wildlife habitat (Bencin et al., 2016). Crop-raiding by wildlife can reduce agricultural yields by over 10% (Anderson & Pariela, 2005; Lamarque et al., 2009), thereby jeopardizing local livelihoods and food security. Livestock predation by carnivores also threatens human livelihoods. Losses from depredation on lands surrounding Waza National Park, Cameroon are approximately $220,000 annually (Lamarque et al., 2009). Finally, attacks by wildlife, primarily crocodiles, cause an average of 118 reported deaths per year in Mozambique (Dunham et al., 2010). As retaliation against wildlife and to meet their subsistence needs, humans kill and poach wildlife, degrade wildlife habitat, and overexploit natural resources (Neelakantan et al., 2019). Poaching is fueled by past interactions with wildlife, negative attitudes towards wildlife (Kansky et al., 2014), preemptive efforts to reduce costs associated with living with wildlife, and financial insecurity in rural communities (Kahler & Gore, 2015).

Although the motives for people destroying wildlife and habitat are complex, understanding what drives these behaviors can greatly improve efforts to attain wildlife conservation outside protected areas. People’s attitudes and perceptions affect their behavioral intentions (Ajzen, 1991) and inform the cultural and socio-political context of wildlife conservation. People’s tolerance for wildlife and individual species are good predictors of potential retaliatory behaviors such as poaching, poisoning, or habitat degradation (Kahler & Gore, 2015). As such, understanding people’s tolerance for wildlife is necessary for managers to better design policies aimed at reducing human-wildlife conflicts (Kansky & Knight, 2014; Mir et al., 2015).

It was previously assumed that tolerance for wildlife is predominantly shaped by the costs of living with wildlife, but recent research suggests that this assumption is an oversimplification (Dickman et al., 2014; Kansky et al., 2016; Bencin et al., 2016). Unfortunately, conservationists have designed policies and programs based on the assumption that people’s response to human-wildlife conflicts is directly proportional to the amount and frequency of wildlife damages, and that reducing damages increase support for wildlife conservation (Dickman et al., 2014). Accordingly, conservationists have focused on reducing crop destruction, livestock predation, and human injury and death by wildlife. However, these interventions are not always effective (Eklund et al., 2017) and can increase intangible costs such as reduced school attendance by children to guard livestock, time and money spent on mitigation efforts, and loss of sleep worrying about problem animals (Ogra, 2008). In a meta-analysis of attitudes toward conflict species, Kansky et al., (2014) found that intangible costs were the best predictors of attitudes. Additional studies have found that socio-demographics (gender, age, education, ethnicity, religion, wealth, and length of residence in an area), potential exposure to human-wildlife conflicts (livestock holdings and depredation, crop damage, distance between communities and protected areas, and restricted access to resources owing to the creation of protected areas), knowledge of wildlife, and people’s level of agreement with wildlife management decisions influence attitudes toward wildlife and protected areas (Karlsson & Sjöström, 2007; Ogra, 2008; Shibia, 2010; Guerbois et al., 2013; Dickman et al., 2014; Mir et al., 2015; Bencin et al., 2016; Mkonyi et al., 2017; Ntuli et al., 2019). Socio-demographics may positively or negatively affect people’s tolerance of wildlife, and these impacts may vary regionally (Mir et al., 2015). Regional differences in people’s attitudes towards wildlife and their willingness to coexist with wildlife highlight the importance of conducting research on human-wildlife conflicts in under-examined regions of the world.

Relatively little research on human-wildlife conflicts has been conducted in Mozambique. Kansky et al.’s (2014) meta-analysis included no case studies from Mozambique and Browne-Nuñez and Jonker’s (2008) meta-analysis of determinants of human-wildlife conflicts only included a single study from Mozambique. Because the formation of attitudes towards wildlife varies regionally and Mozambique has a unique history of Portuguese rule, livelihood instability from extended conflict, and recent rewilding efforts, findings from surrounding countries may not be generalizable to wildlife conservation efforts in Mozambique. To address this research and policy gap, we implemented a study to examine how people’s tolerance for wildlife in the buffer zones surrounding protected areas in Mozambique are affected by socio-demographic, spatial, and wildlife-related factors.

Our research questions were: (1) which variables are correlated with tolerance for wildlife on community-owned lands surrounding protected areas; and (2) how does tolerance vary spatially? We predicted that:

- Females would be less likely to tolerate wildlife since they are often disproportionately affected by human-wildlife conflict (Ogra, 2008). Household income would be positively correlated with tolerance for wildlife as even relatively small losses can have a major impact on the lives and livelihoods of poorer households.
- Individuals with higher education would be more likely to tolerate wildlife.
- Livestock owners would be less likely to tolerate wildlife because they face the risk of predation.
- Wildlife-related costs (livestock loss, crop damage, and human harm) would reduce tolerance for wildlife.
- People who live farther from the protected area would be more likely to tolerate wildlife because they would be less likely to encounter wildlife, experience human-wildlife conflicts, or come into conflict with park staff over the management of wildlife.
- People’s tolerance for wildlife would be reduced by restricted access to resources based on the creation of the protected area.
- People who agree with rules and regulations governing wildlife would be more tolerant of wildlife.
- People who received wildlife-based benefits would be more tolerant of wildlife.

## Study area

We conducted this study in Mangalane, an approximately 50,000 ha communal area bordering Sabie Game Park (referred to as Sabie) in southwestern Mozambique (see Fig 1). Mangalane consists of approximately 480 households and 1800 residents who rely primarily on agriculture and livestock for their subsistence needs (Vundla, 2019). In 2000, Sabie was granted a 99-year lease that allowed them to commence operations as a hunting reserve in 2009. Sabie is 28,000 ha and separated from communal lands by approximately 40 km of electric fencing. This fence is designed to minimize human-wildlife conflicts by reducing wildlife’s access to the communal areas and people’s access to the park (Merz, 2014).

**Fig. 1.**
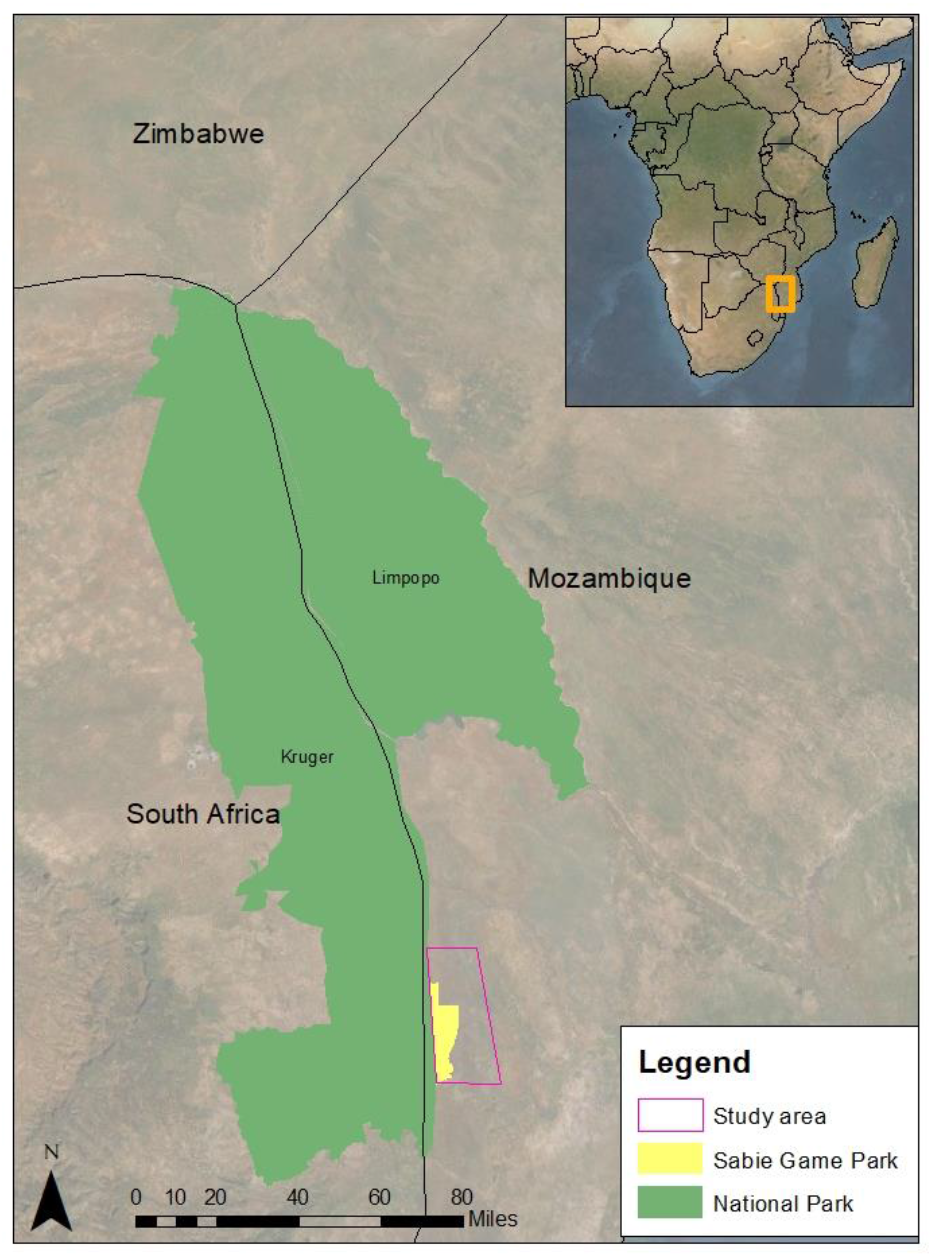
Map of Study Area (outlined in pink) with Sabie Game Park (in yellow) and neighbouring national parks (in green).

Sabie Game Park is part of the Great Limpopo Trans-Frontier Conservation Area, also referred to as a “peace park”. The Great Limpopo Trans-Frotneir Conservation Area includes Kruger National Park in South Africa, Limpopo National Park in Mozambique, and Gonarezhou National Park in Zimbabwe as well as private parks and non-protected areas bordering these parks. The aim is to include local people in the management and economic benefits of biodiversity conservation (Spierenburg et al., 2008).

In 2013 a community-based natural resource management project was initiated in Mangalane to reduce poverty and increase community engagement in wildlife conservation. The project centered on distributing 20 percent of the trophy hunting fees from Sabie to the local community, as mandated by Mozambican law (Merz, 2014; Vundla, 2019). Other initiatives related to this project include environmental education, provision of wells, and human-wildlife conflict mitigation.

## Methods

As part of the community-based natural resource management project, in 2017 Peace Parks Foundation conducted a household livelihood and attitudinal survey in communities bordering Sabie. The survey included questions on attitudes, demographics, health, education, natural resource use, and livelihoods. A total of 237 individuals were interviewed and participation was completely voluntary. Individuals were purposively selected according to a convenience sample from those individuals that were home at the targeted households at the time of sampling. The survey was conducted orally in Shangaan by trained interviewers from the community. Of the 237 surveys, we removed 12 incomplete surveys and 1 extreme spatial outlier, leaving 224 complete surveys used for analysis (94.5% completion rate). We used this survey data to analyze community member’s tolerance for wildlife.

We used the survey question “Do you think it is important to have wildlife in your area?” (yes=1, no=0) as an indicator of wildlife tolerance. The term used for wildlife refers to any wild and non-domesticated animals; we did not differentiate between species or types of wildlife. We analyzed responses to this question using a generalized logistic regression on all possible combinations of explanatory variables using R version 3.5.0 and the MuMin package (R Core Team, n.d.; Barton, 2009). We included 15 explanatory variables in the logistic regression models: gender; age; education; employment (a proxy for household income); cattle ownership; goat ownership; crop loss; livestock predation; human harm by wildlife; restricted access to natural resources owing to the creation of Sabie; agreement with the rules governing wildlife conservation; receipt of wildlife benefits and the value of wildlife-based benefits received; and distances between the community and the park fence and park gate respectively (see Table 1). We also included interaction variables in the estimated models. We considered coefficients to be statistically significant at the 0.1 level. We selected the best-fit model based on the lowest Akaike Information Criteria (AIC) value.

**Table 1.**
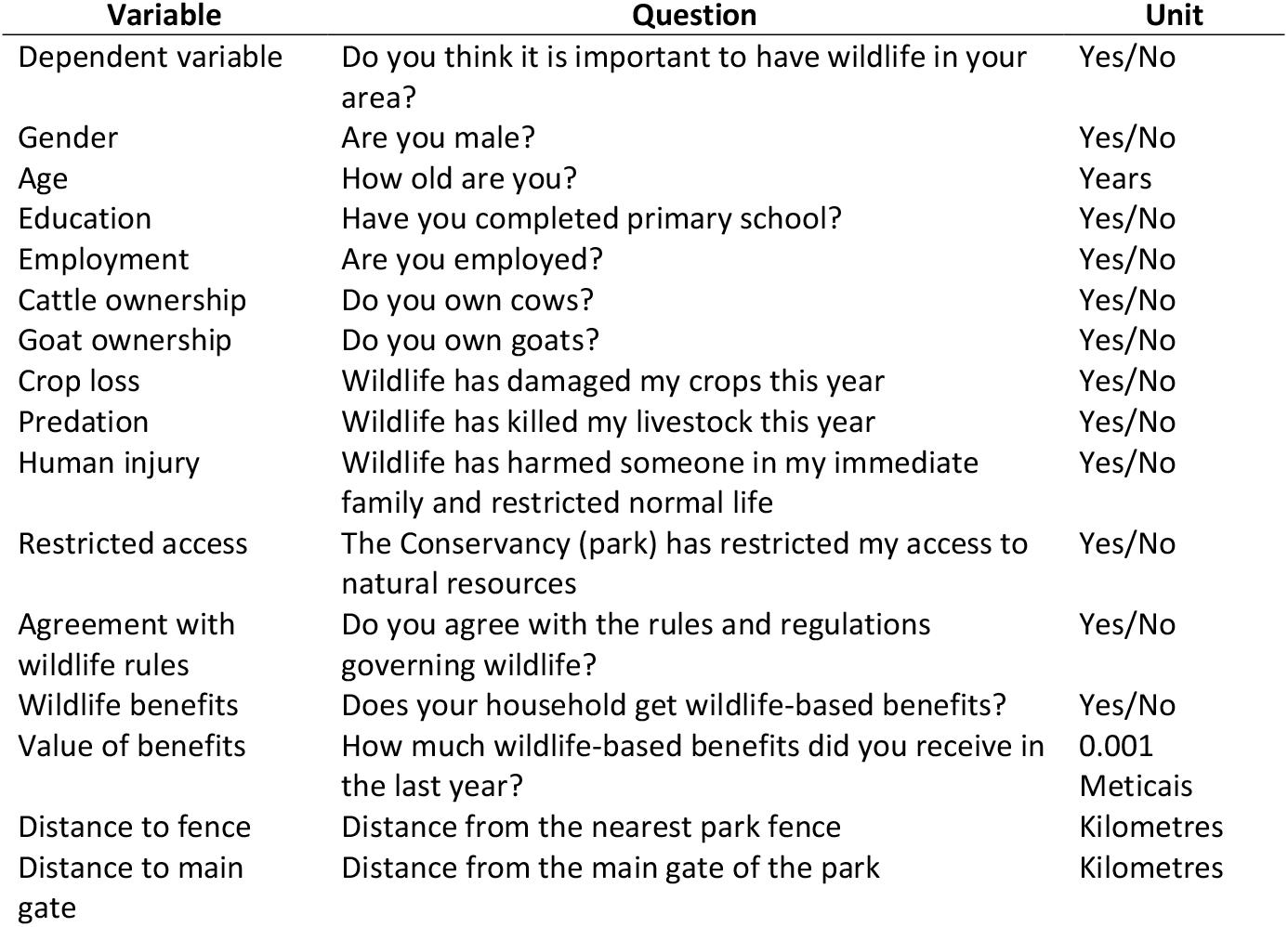
Survey questions used to derive the dependent variable and 15 predictor variables used in logistic regression analysis.

Each of the surveys included in our analysis contained answers to the wildlife tolerance question and the enumerators had recorded the GPS coordinates for the houses of these respondents. We used these data to test for spatial autocorrelation and analyze the spatial distribution of tolerance for wildlife. We tested for global spatial autocorrelation using a global Moran’s I. We conducted a Getis-Ord hot-spot analysis (Gi*) to identify areas that had higher or lower tolerance than expected by random selection (Ord & Getis, 1995). We used the optimization method in ArcMap to determine the distance for calculation (ESRI, 2013) because there were no specific boundaries for this area. The optimization used the average distance to a respondent’s 11 nearest neighbors (1.76 km).

## Results

Of the 224 participants included in this analysis, the majority were tolerant of wildlife (n=137). Most participants agreed with wildlife rules, had lost crops to wildlife in the previous year, had restricted access to natural resources, experienced livestock predation in the previous year, had some education, were female, were unemployed, and did not receive wildlife benefits (see Table 2). The mean age of respondents was 43.0 years (SD=19.7 years). The mean level of benefits received was 428.35 Meticais or $7.26 (SD=1520.54 Meticais). Respondents lived an average of 5.88 km (SD=3.85 km) from the nearest park fence and 20.85 km (SD=11.47 km) from the main gate to the park.

**Table 2.** Aggregate data for binary variables separated by wildlife tolerance (yes and no categories refer to the response to the binary predictor variable question).

| Binary Variables | Aggregate Data |  |  |  | Respondents who are tolerant of wildlife |  |  |  | Respondents who are intolerant to wildlife |  |  |  |
| --- | --- | --- | --- | --- | --- | --- | --- | --- | --- | --- | --- | --- |
|  | Yes |  | No |  | Yes |  | No |  | Yes |  | No |  |
|  | number | percent | number | percent | number | percent | number | percent | number | percent | number | percent |
| Agreement with rules | 183 | 82% | 41 | 18% | 123 | 90% | 14 | 10% | 60 | 69% | 27 | 31% |
| Crop loss | 173 | 77% | 51 | 23% | 116 | 85% | 21 | 15% | 57 | 66% | 30 | 34% |
| Restricted access | 160 | 71% | 64 | 29% | 114 | 83% | 23 | 17% | 46 | 53% | 41 | 47% |
| Predation | 132 | 59% | 92 | 41% | 93 | 68% | 44 | 32% | 39 | 45% | 48 | 55% |
| Education | 128 | 57% | 96 | 43% | 74 | 54% | 63 | 46% | 54 | 62% | 33 | 38% |
| Goat ownership | 104 | 46% | 120 | 54% | 64 | 47% | 73 | 53% | 40 | 46% | 47 | 54% |
| Human injury | 99 | 44% | 125 | 56% | 67 | 49% | 70 | 51% | 32 | 37% | 55 | 63% |
| Cattle ownership | 98 | 44% | 126 | 56% | 60 | 44% | 77 | 56% | 38 | 44% | 49 | 56% |
| Male | 94 | 42% | 130 | 58% | 61 | 45% | 76 | 55% | 33 | 38% | 54 | 62% |
| Wildlife benefits | 59 | 26% | 165 | 74% | 46 | 34% | 91 | 66% | 13 | 15% | 74 | 85% |
| Employment | 49 | 22% | 175 | 78% | 27 | 20% | 110 | 80% | 22 | 25% | 65 | 75% |

The majority of respondents who agreed with the rules governing wildlife were tolerant of wildlife (see Table 2 and Figure 3). Surprisingly, the majority of respondents who experienced restricted access to natural resources owing to the creation of the protected area, crop loss from wildlife, and livestock predation were tolerant of wildlife. More respondents who had received wildlife benefits were tolerant of wildlife than intolerant, although only 59 respondents had received any wildlife benefits. A higher proportion of men were tolerant of wildlife whereas a higher proportion of females were not tolerant of wildlife. The mean age of respondents who tolerated wildlife was 44.1 years (SD=20.6 years), slightly higher than the mean age of 41.3 years (SD=17.9 years) for respondents who did not tolerate wildlife. Respondents that were tolerant of wildlife lived an average 21.5 km (SD=11.4 km) from the park fence and 6.5 km (SD=3.9 km) from the park gate while respondents that were intolerant lived an average of 19.9 km (SD=11.5 km) and 4.9 km (SD=3.5 km) from the fence and gate, respectively. Respondents that tolerated wildlife received an average of 419.07 Meticais or $7.10 (SD=1250.36 Meticais) per year in wildlife benefits, compared to an average of 442.99 Meticais or $7.51 (SD=1868.30 Meticais) per year in benefits received by respondents that were not tolerant of wildlife. The comparative trends in continuous variables by wildlife tolerance is shown in Fig 4.

Because wildlife tolerance depends on a combination of respondents’ previous experiences with wildlife, identities, and values, regression analysis is required to illuminate which factors are correlated with tolerance for wildlife, taking all other respondent characteristics into account. We used logistic regression analysis to analyse which combination of respondent characteristics were correlated with tolerance for wildlife. We found that gender, age, agreement with wildlife rules, receipt of wildlife benefits, distance to fence, and restricted access to resources best predicted whether respondents were tolerant of wildlife (see Table 3). Men, older respondents, respondents who lived farther from the Sabie fence, and respondents whose access to natural resources had been restricted by the creation of Sabie were likely to be more tolerant of wildlife, all else being held equal. Respondents whose households had received wildlife-based benefits were approximately 4 times more likely to be tolerant of wildlife (odds ratio=4.129), all else being held equal. Although respondents who agreed with the rules governing wildlife were more likely to be tolerant of wildlife, this effect was offset by the age of respondents (negative coefficient on the interaction between agreement with wildlife rules and the age of the respondent). Respondents’ agreement with wildlife rules had the greatest impact on their tolerance for wildlife, with individuals who agreed with these rules being approximately 30 times more likely to tolerate wildlife (odds ratio=30.240), all else being held equal. However, this effect disappeared for respondents who were over 86 years of age. All other variables that we predicted would be correlated with tolerance for wildlife (employment, livestock ownership, human harm, crop loss, predation, distance to park gate, and education) were not statistically significant and were omitted from the best-fit model. Additional interaction terms included in the model were also not statistically significant and consequently were excluded from the best-fit model.

**Table 3.**
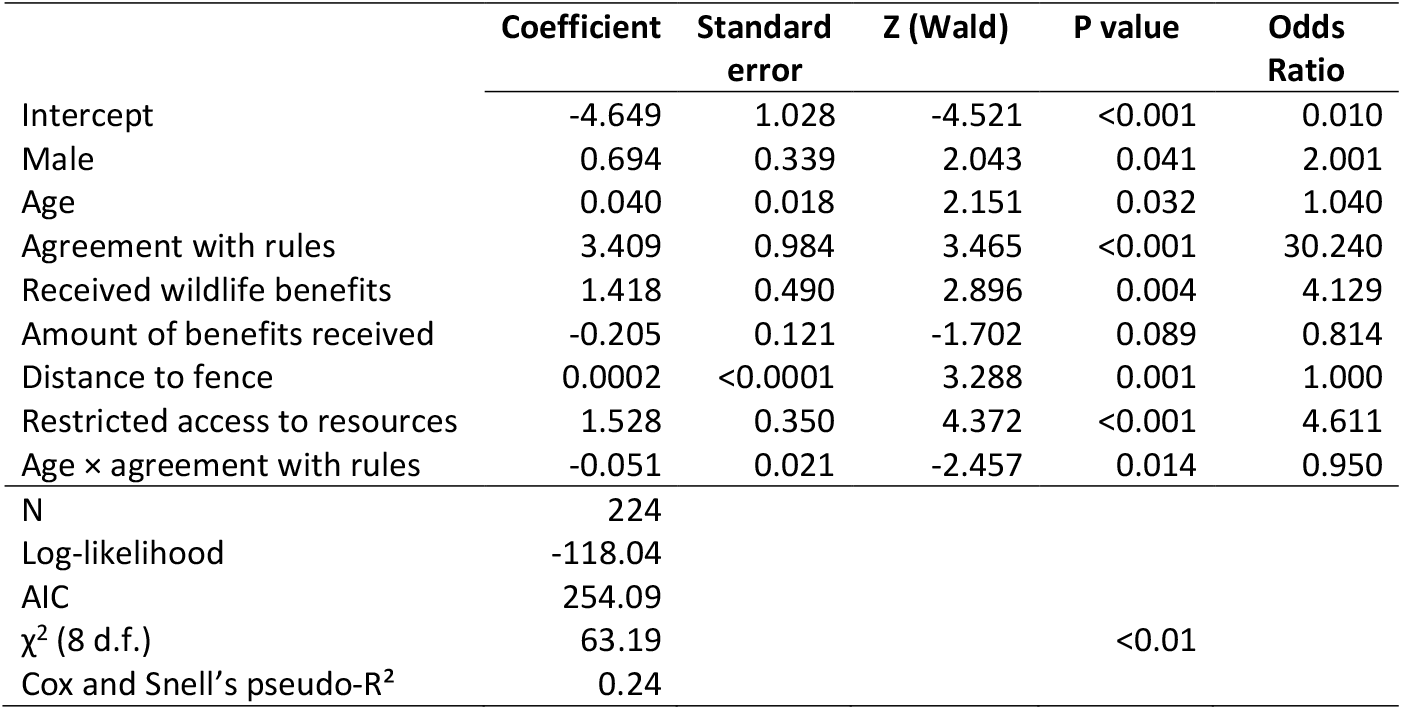
Results of the best-fit logistic regression model according to the lowest Akaike Information Criteria (AIC) value of all possible models run including relevant interaction terms.

|  | **Coefficient** | **Standard error** | **Z (Wald)** | **P value** | **Odds Ratio** |
| --- | --- | --- | --- | --- | --- |
| Intercept | -4.649 | 1.028 | -4.521 | <0.001 | 0.010 |
| Male | 0.694 | 0.339 | 2.043 | 0.041 | 2.001 |
| Age | 0.040 | 0.018 | 2.151 | 0.032 | 1.040 |
| Agreement with rules | 3.409 | 0.984 | 3.465 | <0.001 | 30.240 |
| Received wildlife benefits | 1.418 | 0.490 | 2.896 | 0.004 | 4.129 |
| Amount of benefits received | -0.205 | 0.121 | -1.702 | 0.089 | 0.814 |
| Distance to fence | 0.0002 | <0.0001 | 3.288 | 0.001 | 1.000 |
| Restricted access to resources | 1.528 | 0.350 | 4.372 | <0.001 | 4.611 |
| Age × agreement with rules | -0.051 | 0.021 | -2.457 | 0.014 | 0.950 |
| N | 224 |  |  |  |  |
| Log-likelihood | -118.04 |  |  |  |  |
| AIC | 254.09 |  |  |  |  |
| $\chi^2$ (8 d.f.) | 63.19 | | | <0.01 | |
| Cox and Snell's pseudo-R<sup>2</sup> | 0.24 |  |  |  |  |

Because distance to Sabie was included in the best-fit model, we tested for global and local spatial autocorrelation to determine if there was a spatial pattern of clustering of wildlife tolerance. Figure 2 shows the distribution of respondents by wildlife tolerance. The spatial analyses showed no global autocorrelation with a Moran’s I of 0.065 (p=0.637). The Gi* hotspot analysis at an optimized 1.76 km also revealed no statistically significant (p<0.05) hot or cold spots in tolerance for wildlife. Therefore, we did not pursue additional spatial analyses of wildlife tolerance.

**Fig. 2.**
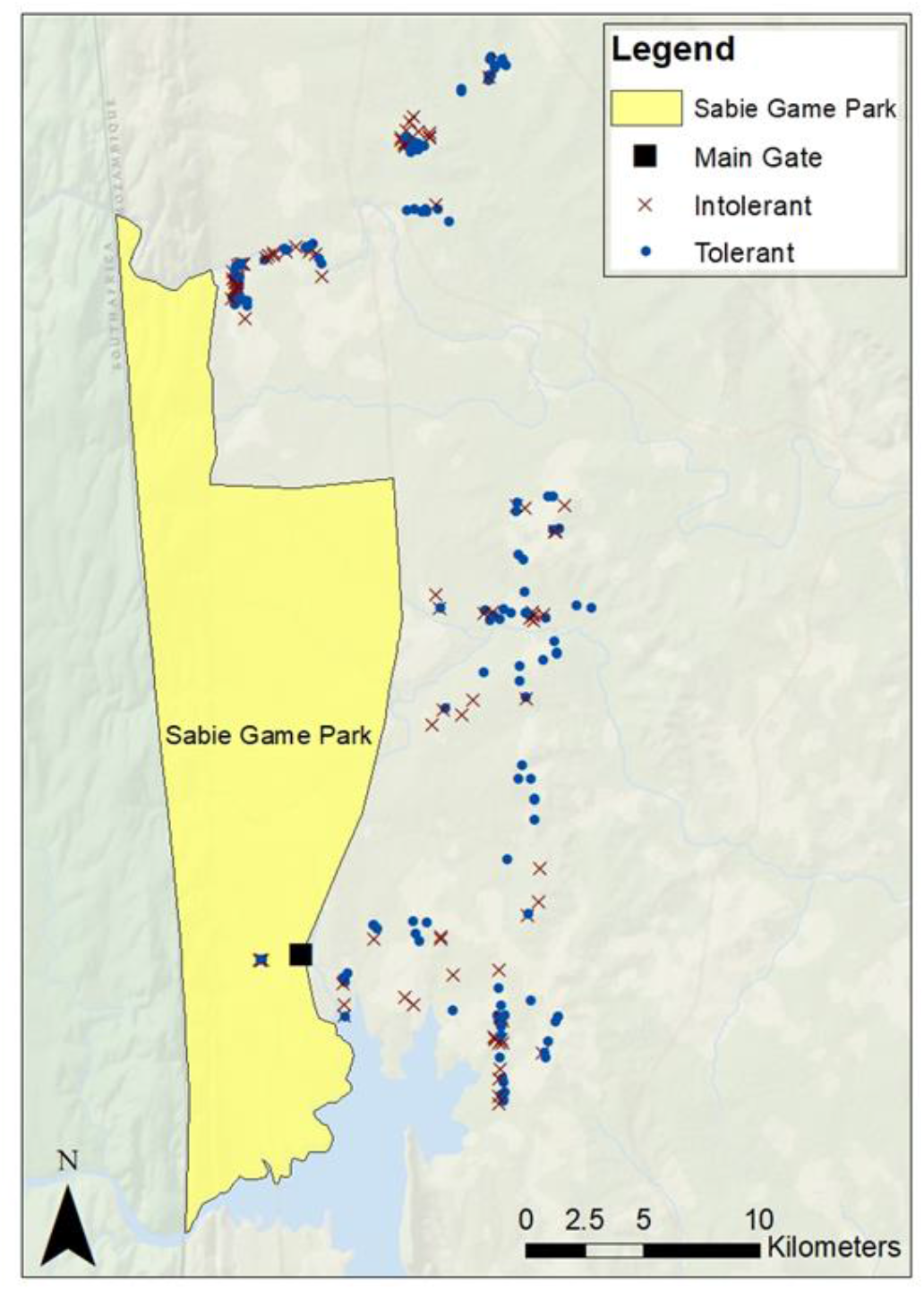
Spatial Distribution of households surveyed according to wildlife tolerance with tolerant respondents shown with a blue dot and intolerant respondents shown with a red x.

**Fig. 3.**
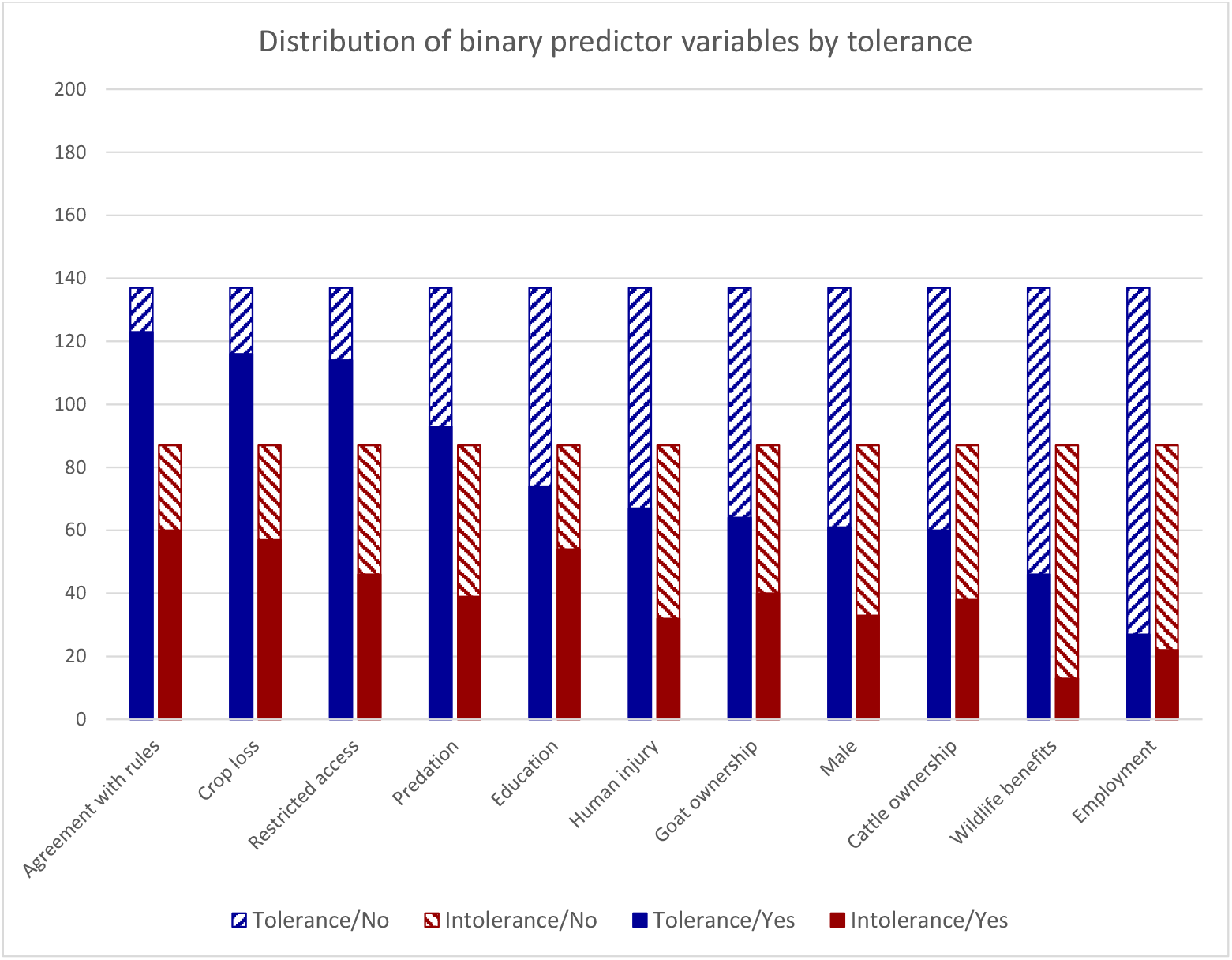
Distribution of binary predictor variables (employment, wildlife benefits, gender, cattle ownership, human injury, goat ownership, education, predation, restricted access, crop loss, and agreement with rules) by wildlife tolerance. Individuals that were tolerant of wildlife are displayed in blue (n=137) and individuals that were intolerant of wildlife are displayed in red (n=87). Tolerance is sub-divided by response to the predictor variable question with yes displayed with a solid colour and no with a hatched pattern.

**Fig. 4.**
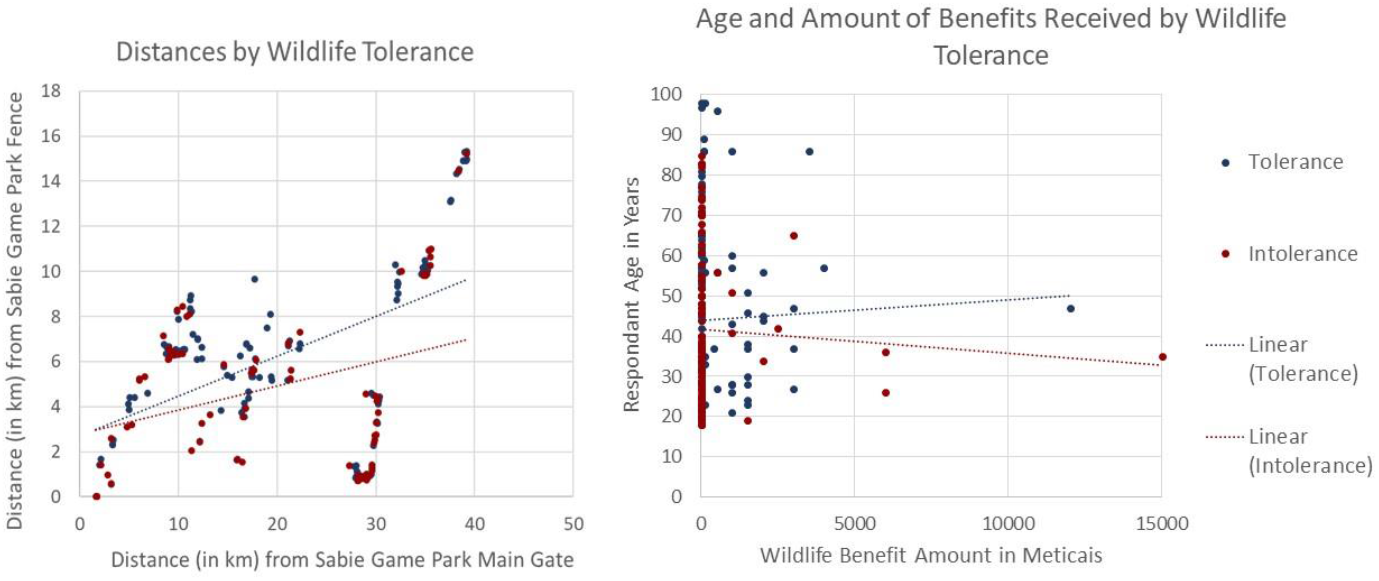
Scatterplot comparison of continuous predictor variables (household distance from the Sabie Game Park main gate by household distance from Sabie Game Park fence on the left and age by amount of benefits received on the right). Wildlife tolerance is displayed in blue and intolerance in red. Linear trend lines for tolerance and intolerance are included as dotted blue lines for tolerance and dotted red lines for intolerance.

## Discussion

Biodiversity conservation relies on land outside of protected areas where wildlife often come into conflict with humans. Understanding the drivers of tolerance is vital for ensuring the coexistence of people and wildlife in these shared spaces. We found that community members in Mangalane tended to be tolerant of wildlife and that tolerance for wildlife did not exhibit spatial patterns. Our findings corroborate more recent literature that tolerance for wildlife is not directly and proportionally related to human-wildlife conflicts (Kansky et al., 2014, 2016; Mkonyi et al., 2017; Broekhuis et al., 2020). However, many management efforts continue to focus on mitigation of direct costs as a means of improving conservation despite evidence that these efforts are ineffective. Cattle predation rates are typically low, affecting less than 3% of cattle (Thorn et al., 2012; Dickman et al., 2014). People may, therefore, be willing to tolerate small amounts of predation or crop loss. Low levels of livestock depredation may explain why men, who are generally responsible for cattle, were more likely to tolerate wildlife. As we predicted, agreement with rules governing wildlife and greater distance from Sabie increased tolerance for wildlife. Households that are farther from the park are likely to have fewer interactions with wildlife and therefore fewer opportunities for conflict. Continued efforts to mitigate human-wildlife conflicts that appear to be considered tolerable by community members are a poor use of resources.

Contrary to our predictions, restricted access to natural resources associated with the creation of Sabie increased tolerance for wildlife. Prior studies in Zimbabwe (Guerbois et al., 2013) and Kenya (Shibia, 2010) showed that resentment over not being allowed to sustainably utilize natural resources in protected areas led to negative attitudes toward parks and wildlife. In contrast to these studies, Sabie is surrounded by a game fence that effectively keeps most wildlife away from communities. Although the fence restricts access to resources, it may be perceived as a benefit because it keeps human-wildlife conflicts at a tolerable level. However, allowing access to protected areas in drought years may be important to maintain tolerance for wildlife (Hazzah et al., 2013). Access restrictions in Mangalane may become more costly in drought years if there is insufficient water and vegetation outside Sabie to support livestock.

In contrast to Kansky et al. (2014), we found that if respondents received wildlife-based benefits they were more likely to be tolerant of wildlife. On average, respondents received a relatively small amount of $7.26 (428.4 MT) per year in benefits (maximum of $254 or 15,000 MT). However, our findings also suggested that as benefits increase tolerance for wildlife may decrease, i.e., the receipt of some benefits is important, but payments should not be set too high (although this was only significant at the 0.1 level). Providing payments may create a market mentality that results in self-interested behavior (Reeson & Tisdell, 2006), especially if rewards are not appropriately linked to desired behavior such as wildlife conservation (Lepper & Greene, 2015). The equitable distribution of benefits may be more important in promoting tolerance for wildlife (Groom & Harris, 2008). Additional research in Mangalane is needed to determine the appropriate amount and type (e.g., cash versus in-kind) of benefits needed to maintain tolerance for wildlife. If relatively small amounts of benefits distributed evenly can significantly improve tolerance, as indicated in these results, this may be a cost-effective means of attaining desirable wildlife conservation outcomes.

Although our results provide important insights into factors influencing tolerance for wildlife in Mozambique, the survey was not specifically designed for this purpose. This survey only asked if households had experienced wildlife-based damages in the past year, but tolerance can be influenced by historic events and the degree of damages, neither of which we captured in this study. We used employment as a proxy for wealth, but a more accurate measure of household income may provide a better understanding of how wealth influences tolerance for wildlife. Further research is required to augment and validate our findings.

Nonetheless, our study reinforces recent findings that tolerance for wildlife is not simply determined by human-wildlife conflicts. Continuing to operate under broad assumptions that reducing human-wildlife conflicts and increasing wildlife-based benefits will yield greater tolerance for wildlife and appropriate conservation behaviors by community members is counterproductive. Equitable distribution of wildlife-based benefits and efforts to include community members in wildlife conservation decision-making so that they recognize the legitimacy of rules related to wildlife may be more productive.

Mozambique has been under-represented in studies of wildlife tolerance in Africa. With a unique history of Portuguese colonization and extended civil war, it is difficult to generalize results from other parts of sub-Saharan Africa to Mozambique without assessing them in the local context. This study provides valuable evidence of the strength of previous findings far beyond their local contexts. Because these results from Mozambique corroborate regional trends in wildlife tolerance, we provided further evidence that recent recommendations on how to improve wildlife tolerance may be applied more broadly across sub-Saharan Africa. Efforts to provide equitable wildlife-based benefits should be prioritized over reducing human-wildlife conflicts. Establishing buffer zones outside of protected areas for sustainable resource use, but not housing, may further reinforce tolerance for wildlife. Involving local communities in decision making may improve agreement with conservation rules which was linked to increased tolerance for wildlife in our study. Conservation efforts should include women to increase their tolerance for wildlife. Community-based natural resource management projects that allow community participation in decision-making processes and promote equitable benefit sharing should promote tolerance for wildlife and wildlife conservation behaviors at the community level.

## Author contributions

Study design: LM and TF; fieldwork: SM; data analysis: LM; writing the article: LM and EP; Editing the article: LM, EP, TF, and SM.

## Acknowledgements

We would like to thank Peace Parks Foundation for their work in conducting the household livelihood survey data and their willingness to share the dataset. We are extremely grateful to the 237 anonymous individuals who dedicated time to participate in the survey. Thank you to the data collectors from Mangalane: Aauroia, Amos, Bongi, George, Jeremiah, Louis, Ivonne, and Zitu. We appreciate Caroline Huguenin for helping with preliminary analysis, James Martin for helping with R coding, and Dr. Jane Southworth for her valuable feedback and encouragement. Finally, we are grateful for the helpful comments from anonymous reviewers.

## Conflicts of interest

None

## Ethical standards

Peace Parks Foundation is a registered non-profit organization operating in Mozambique. They work with government agencies to manage the Great Limpopo Trans-frontier conservation area and other protected areas throughout Southern Africa. They undertook the research used in this study for their own purposes in 2017 and kindly offered the use of their dataset for this research in 2019. While Mozambique does not have a formal IRB process for non-medical research on human-subjects, Peace Parks complied with all local and national standards for research. This included obtaining permission from Chief Mangalane, Chef de Poste, and private game park owners who helped select the group of nine local facilitators who helped with data collection. Survey participation was voluntary, and each participant provided oral consent. Author, Shylock Muyengwa was contracted by Peace Parks Foundation to conduct the survey research used in this study.

## Notes

### Competing Interest Statement

The authors have declared no competing interest.

